# Habit and the hippocampus: Model-based spatial representations without outcome-sensitive control

**DOI:** 10.64898/2026.03.06.710211

**Authors:** Shiyun Wang, Ryan Grgurich, Hugh T. Blair

**Affiliations:** UCLA Psychology Department, 90095

## Abstract

Goal-directed decisions depend on predictions of future outcomes and are thus sensitive to revaluation of expected outcomes, whereas habitual decisions depend on cached action values or stimulus-response associations and are insensitive to outcome revaluation. Hippocampal neurons encode future states (potentially supporting prediction of future outcomes during goal-directed choice) as well as present states (potentially supporting retrieval of state-dependent action values during habitual choice). Prior studies suggest a role for the hippocampus in goal-directed, outcome-sensitive decision making, but few studies have examined its role in habitual, outcome-insensitive action selection. Here, we tested this using a two-outcome spatial navigation task in which identical turning actions (left or right) led either to preferred milk or non-preferred water outcomes, depending on the spatial location at which the choice was made. Rats (20 males, 22 females) were trained under conditions that either required hippocampal-dependent discrimination between choice points or allowed hippocampal-independent navigation using visual beacons. Rats then underwent 24 h water restriction during which separate cohorts underwent Contingency Updating (CU), in which rats re-experienced the maze action-outcome contingencies while thirsty, or Incentive Learning (IL), in which rats received only a two-bottle preference test while thirsty without re-experiencing maze contingencies. Both cohorts were later tested under thirst on an unbaited maze. CU rats shifted their choices toward water-associated responses, whereas IL rats perseverated on milk-associated responses, even when navigation depended on the hippocampus. These results provide novel evidence that hippocampal-dependent navigational decisions can show insensitivity to outcome revaluation, a widely-used behavioral criterion for habitual control.

**Significance Statement:** We show here that the dorsal hippocampus can support navigational decisions that are insensitive to outcome revaluation during a spatial task in which identical actions lead to different outcomes depending on location. Even when the hippocampus was required for accurate navigation to reach a preferred outcome, rats overtrained on this task failed to adjust their choices after revaluation altered their outcome preferences. To our knowledge, these findings provide the first direct evidence that hippocampal-dependent navigation can satisfy a widely used behavioral criterion for habitual control.

## Introduction

The hippocampus has long been implicated in spatial navigation, relational memory, and model-based planning. During associative learning, hippocampal contributions are especially prominent when predictive stimuli consist of spatial contexts or configural cue patterns (Hirsh, 1974; Sutherland and Rudy, 1989; Eichenbaum et al., 1994; Eichenbaum and Cohen, 2014). In spatial navigation tasks, the hippocampus is critical when behavior depends on internal models or “cognitive maps” of the environment but is not required when tasks can be solved using simple response strategies or cue-response associations (Olton and Samuelson, 1976; O’Keefe and Nadel, 1978; Morris et al., 1982; Packard and McGaugh, 1992, 1996; McDonald and White, 1994; Oliveira et al., 1997).

Parallel distinctions have emerged in studies of goal-directed versus habitual decision making. Goal-directed actions are operationally defined by their sensitivity to outcome revaluation and are thought to rely on mental models, whereas habitual actions are operationally defined by insensitivity to outcome revaluation and are thought to depend on cached action values or stimulus-response associations (Dickinson, 1985; Daw et al., 2005; Dolan and Dayan, 2013; Balleine, 2019). With extended training, behavior often transitions from outcome-sensitive to outcome-insensitive control, accompanied by a shift from prefrontal and dorsomedial striatal involvement toward dorsolateral striatum (Adams, 1982; Balleine and Dickinson, 1998; Yin and Knowlton, 2006; Balleine and O’Doherty, 2009). A similar transition has been observed in dual-strategy T-maze tasks, where rodents initially use a hippocampal-dependent place strategy and then later adopt a non-hippocampal response strategy with overtraining (Packard and McGaugh, 1992, 1996; White et al., 2013).

Outcome sensitivity has rarely been assessed in spatial navigation tasks. Kosaki et al. (2018) reported evidence that on the dual-strategy T-maze, the place strategy was sensitive to outcome revaluation whereas the response strategy was not. In non-spatial decision tasks, early studies reported that the hippocampus does not influence outcome sensitivity (Corbit and Balleine, 2000; Ward-Robinson et al., 2001; Corbit et al., 2002), but later work showed that outcome revaluation during early stages of lever pressing depended on both the hippocampus and spatial context before eventually becoming independent of both (Bradfield et al., 2020). These findings accord with other lines of evidence that the hippocampus participates in goal-directed decisions when they depend on spatial or contextual information (Chang and Gold, 2004; Miller et al., 2017). Consistent with this view, hippocampal place cells generate “preplay” sequences preceding navigational choices that may support prospective evaluation of future outcomes (Johnson and Redish, 2007; Pfeiffer and Foster, 2013; Mattar and Daw, 2018; Kay et al., 2020), and some hippocampal neurons have recently been hypothesized to encode successor representations that convey policy-weighted predictions of future states (Gershman et al., 2012; Stachenfeld et al., 2017; Garvert et al., 2023; Geerts et al., 2024).

While such findings suggest how hippocampal representations may prospectively encode future states, classical cognitive-map theories emphasized encoding of the animal’s current spatial position rather than future trajectories (O’Keefe and Dostrovsky, 1971; O’Keefe and Nadel, 1978). Reinforcement learning theories posit that cached action values are attached to current task states, so if hippocampal representations encode present states, then these representations could support habitual action selection. However, few studies have examined the role of hippocampal representations in habit-driven decision making.

Here, we addressed this question using a two-outcome spatial decision task in which identical turning actions led to different rewards (milk or water) depending on choice-point location. Rats were trained under conditions that either required hippocampal-dependent discrimination between choice locations or allowed hippocampal-independent performance using visual beacons, as confirmed by intrahippocampal muscimol inactivation. This design allowed us to determine whether hippocampal-dependent navigation can occur under conditions in which navigational choices are insensitive to outcome revaluation.

## Materials and Methods

All experimental procedures were approved by the Chancellor’s Animal Research Committee of the University of California, Los Angeles, in accordance with the US National Institutes of Health (NIH) guidelines, protocol #2017-038.

### Subjects

Forty-two Long–Evans rats (22 females, 20 males; 8–12 weeks old at arrival; Charles River Laboratories) were used in the experiment. Animals were singly housed in a temperature-and humidity-controlled vivarium under a 12-hour reverse light cycle (lights off at 7:00 AM). All procedures were conducted in accordance with institutional and NIH guidelines for the care and use of laboratory animals.

Seven days after arrival, rats underwent stereotaxic surgery for bilateral dorsal hippocampal cannula implantation (see Surgical Procedures below), except where noted for specific control groups. Following a 7-day recovery period, animals were gradually reduced to 80% of their ad libitum body weight via controlled daily feeding. During this period, rats received handling, reward familiarization, apparatus exposure, and initial training on the corner maze task (see Behavioral Procedures below).

As shown in Table 1, animals were assigned to one of three task conditions: Path Integration (PathInt), Texture, or Beacon. Four male rats assigned to the PathInt condition failed to acquire the task after 40 training days and were dropped from the study. Within each task condition, rats were further assigned to either a Contingency Updating (CU) group or an Incentive Learning (IL) group.

**Table 1.** Behavioral analyses cohort sizes. Breakdown of number and sexes of subjects included in the behavior analyses of the six experimental groups.

|  | PathInt | Texture | Beacon |
| --- | --- | --- | --- |
| Contingency Updating | 3 (2F, 1M) | 6 (3F, 3M) | 11 (7F, 4M) |
| Incentive Learning | 7 (4F, 3M) | 6 (3F, 3M) | 5 (3F, 2M) |

*Note: in the Beacon-CU group, 5 (2F, 3M) rats were not implanted with cannula, and 6 (5F, 1M) received aCSF infusion on the contingency updating day*
*Note: the four male rats that failed to acquire the PathInt task were excluded from this table and from behavior analyses.*

Subject counts included in infusion analyses are shown in Table 2. Three PathInt-IL rats were excluded for having incorrect injector cannula placement.

**Table 2.** Infusion analyses cohort sizes. Breakdown of number and sexes of subjects included in the infusion analyses of the three experimental groups.

|  | PathInt | Texture | Beacon |
| --- | --- | --- | --- |
| Incentive Learning | 4 (2F, 2M) | 6 (3F, 3M) | 5 (3F, 2M) |

### Apparatus

Behavioral testing was conducted in a custom-built maze consisting of a 2 m × 2 m square perimeter track surrounding a central plus maze, forming the shape of a 田 (Figure 1A, Movie 1). Each of the four outer corners contained a metal reward well connected via tubing to a programmable syringe pump which delivered undiluted unsweetened condensed milk (UCM, 100 μL) or water (300 μL). Four unique floor textures were placed inside of the reward zones (ceramic tile samples; Home Depot). Texture–zone pairings were fixed per animal but randomized across animals. To minimize odor cues, a non-accessible reward cup containing UCM was placed at each corner (regardless of whether milk or water was delivered there). Sixteen automated transparent acrylic doors, controlled by stepper motors, were positioned between maze segments and could be raised or lowered to dynamically alter maze configuration during testing. Video monitors (24-inch) were mounted above the midpoint of each straight arm of the square perimeter and angled approximately 45° toward the maze surface.

**Figure 1.**
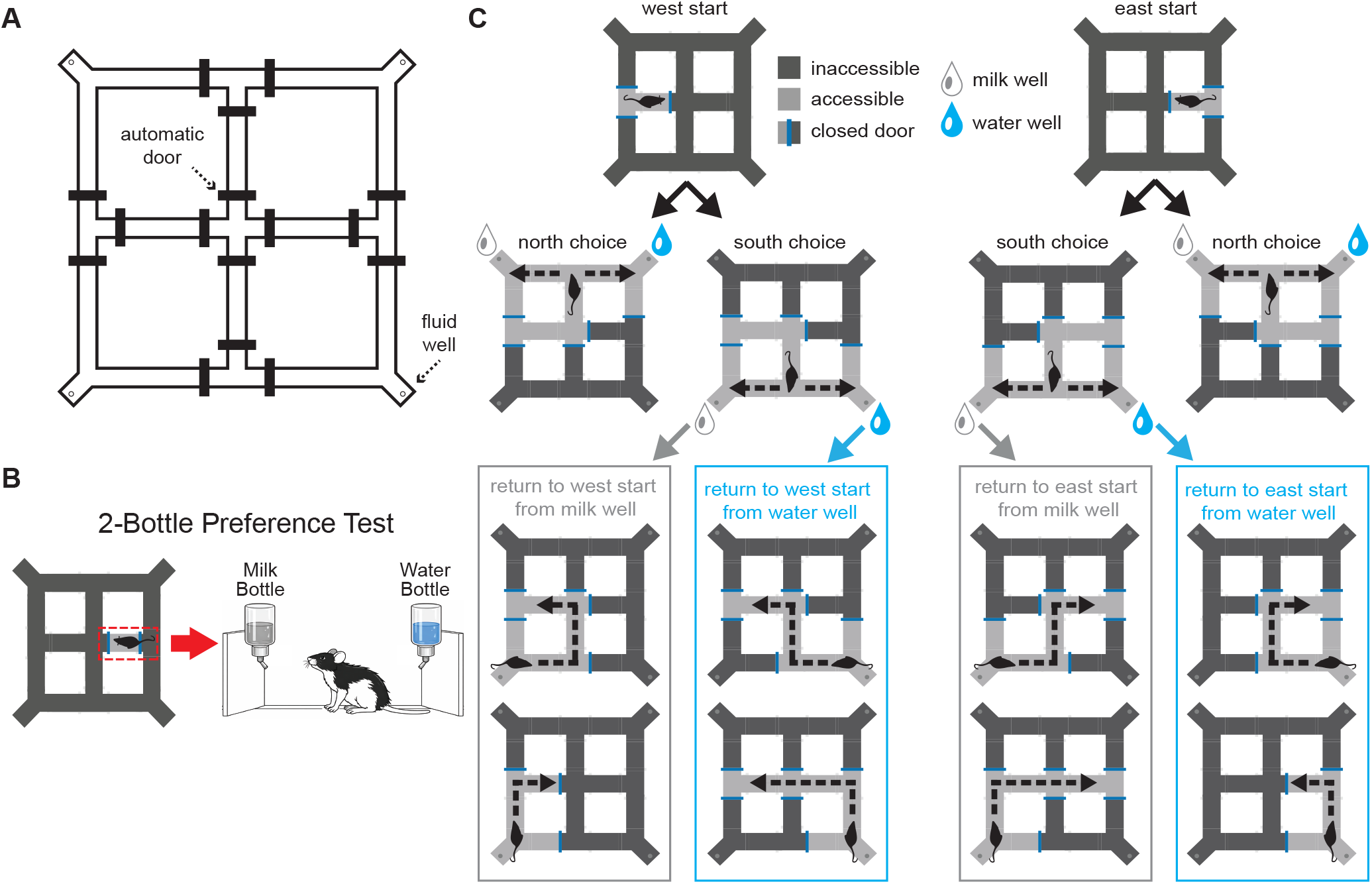
Maze apparatus and task design. A) Overhead diagram of the 2×2 m maze apparatus. B) 2-bottle preference tests were given while rats were confined to the east segment of the maze. C) Task Design: Rat began in the west or east start arm (top row) and was then forced to a north or south choice point (middle row). From the north choice point, left turn leads to milk and right turn leads to water; from the south choice point, the opposite is true. Upon visiting the chosen fluid well, doors were reconfigured to force the rat along one of four return paths to the next start arm (randomly chosen). Bottom row shows return paths from south wells; paths from north wells are identical under vertical reflection (not shown).

All hardware components (including automated doors, syringe pumps, monitors, infrared illumination, speakers, and camera) were controlled via custom software written in Python. The software performed real-time markerless tracking of the rat’s position (30 Hz frame rate) using an overhead video camera, and controlled task phase sequencing by dynamically reconfiguring the maze doors contingent upon the animal’s location and behavior.

The maze was elevated 1 m above the floor in a square testing room whose walls and ceiling were covered with uniform black curtains to eliminate extramaze visual cues. The room was illuminated exclusively with infrared light sources positioned at the four upper corners, allowing for video tracking while rats performed the task in the absence of visible light. White noise was delivered continuously from a centrally mounted ceiling speaker during behavioral sessions to mask extraneous sounds. The maze environment was designed to be visually symmetrical and devoid of stable distal cues. The task reference frame was rotated 90° (clockwise or counterclockwise) relative to the room each day to prevent use of stable extramaze cues, and all maze segments (of identical shape) were interchangeable and were randomly rearranged between sessions (excluding the four reward corner locations) to ensure that task performance could not be guided by fixed intramaze landmarks.

### Cannula implant surgery

Rats were anesthetized with isoflurane (5% for induction in oxygen at 2.5 L/min; maintained at 1.5–2.0%) and placed in a stereotaxic apparatus (Harvard Apparatus). Body temperature was maintained with a heating pad throughout the procedure. Thirty minutes prior to surgery, animals received subcutaneous saline (1 mL) for hydration and carprofen (6 mg/kg; 2 mg/mL solution) for analgesia. Following exposure of the skull and leveling between bregma and lambda, bilateral craniotomies were made targeting the dorsal hippocampus (coordinates relative to bregma: AP −3.96 mm; ML ±3.0 mm). Stainless steel guide cannulae (22 gauge; Protech International Inc.) were lowered to 1.0 mm above the CA1 pyramidal layer (DV −2.5 mm from skull surface). Cannulae were secured to the skull using four stainless steel anchor screws (Bioanalytical Systems Inc.) and dental acrylic (Ortho-Jet BCA Powder, black). Hemostasis was achieved during drilling and implantation using sterile cotton swabs and absorbable gelatin sponge (Surgifoam; Johnson & Johnson MedTech). After the dental cement cured, sterile dummy cannulae were inserted to maintain patency and prevent occlusion. A protective cap was placed over the assembly to prevent removal by the animal. Rats were allowed to recover for seven days prior to initiation of food restriction and behavioral training.

### Behavior task

An overview of the experimental timeline is shown in Figure 2A. Prior to training, animals were given a set of pre-training procedures to familiarize them with the apparatus and instruments. Then, they experienced four phases of the experiment: (1) acquisition training and overtraining, (2) revaluation manipulation (Incentive Learning or Contingency Updating), (3) strategy probing (hydrated and thirsty test sessions), and (4) HPC inactivation.

**Figure 2.**
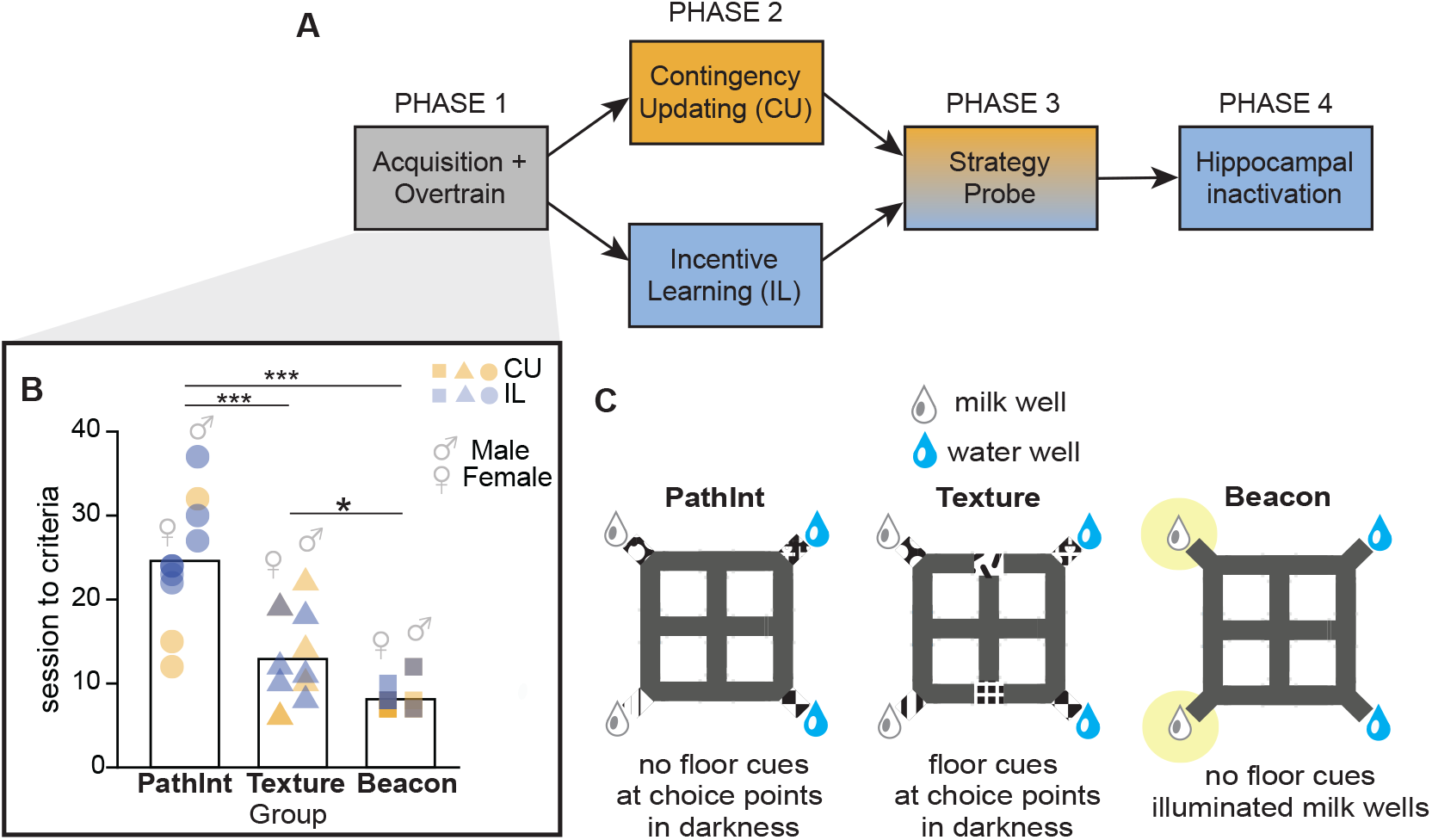
Acquisition and Overtraining A) The experiment consisted of four sequential phases. B) In Phase 1, rats in the PathInt group required more training sessions than Texture or Beacon rats to acquire the task and reach the over-training criterion; a sex difference was observed in the PathInt group (see main text). Planned comparisons: *p < .05, ***p < .001. C) Three groups (PathInt, Texture, Beacon) underwent different training procedures in Phase 1 (see main text).

### Phase 0: Pre-training

#### Handling and Reward Exposure

Animals were handled for 10 minutes daily for four consecutive days. After each handling session, rats were returned to their home cage and were given 10 mL of UCM in a metal cup attached to the cage for 24 hours. Water bottles were replaced with the bottles used in later two-bottle testing to habituate animals to the apparatus. Water consumption was monitored daily.

#### Two-bottle preference test

Rats underwent four days of two-bottle preference testing to establish baseline milk preference. During each session, animals were confined to a straight arm of the maze by raising two barriers (Figure 1B, Movie 2). One bottle contained plain drinking water and the other contained UCM (75 mL each; MidWest Homes for Pets Small Animal Water Bottle). Bottle positions (left/right) were randomized and switched every 4 minutes to control for side bias. Sessions lasted 16–30 minutes on the first day and 8 minutes on subsequent days. Bottles were weighed before and after testing to calculate percent milk consumption (by weight). If a rat consumed <1 g of milk, an additional test was administered the following day.

#### Maze exposure

Rats received two 30-minute free exploration sessions in the maze prior to acquisition training. On both days, reward wells were active and delivered water or UCM upon entry into the reward zones. During the first exposure session, barriers remained static. During the second session, barriers were manually manipulated to allow interaction with different maze configurations. For groups receiving visual cues (see below), the cue was presented for 5 minutes at the start of the second exposure session. The maze was cleaned between sessions and rats using diluted Rescue solution (1:20) for floor segments, 70% ethanol for barriers and textures, and distilled water for reward wells.

### Phase 1: Acquisition Training and Overtraining

All training occurred in darkness under white noise with infrared illumination for tracking.

#### Trial structure

The task is illustrated in Figure 1C and Movie 1. Each trial began with confinement of the rat in one of two start arms (west or east; task reference frame). Access was then granted to one of two spatially distinct choice points (north or south T-intersections). At the choice point, the rat could turn left or right. Once a choice was made, the alternative path was blocked for the remainder of the trial, so that the choice was binary: if choice A was selected on proportion *pA = N_A_/N_total_* of trials, then the alternative choice B was selected on proportion *p_B_ = (1-p_A_)* of trials. Identical turning actions led to different outcomes depending on spatial location. At the north choice point: left → UCM; right → water; at the south choice point: left → water; right → UCM (outcome contingencies were counterbalanced across animals). Thus, successful performance required discrimination of spatial location. The first trial of each session was a forced, unrewarded orientation trial in which the water side was blocked to help animals establish the daily task reference frame. The animal must enter the reward corner it chose on each trial to examine and optionally consume the reward before the intertrial interval is given. One of four intertrial interval return paths (two per start arm) will be randomly chosen and made available. Start arms were randomized across trials. Sessions lasted 30 minutes.

#### Overtraining criterion

Animals advanced to phase 2 after meeting all of the following criteria for two consecutive days: 1) choose milk on ≥8 of the first 10 trials, 2) choose milk on ≥ 90% of all visits to each choice point during the session, and 3) run ≥ 30 trials per session.

#### Mock infusion training

Before each session, animals underwent a mock infusion procedure identical to the real infusion protocol, except injectors were not inserted into the guide cannula. Animals then remained in a holding cage under a black cloth for 25 minutes prior to testing. This was to ensure that rats would remain calm and tolerant of the actual infusion procedure on days when muscimol was delivered.

### Group-specific task variants (Figure 2C)

#### Path Integration (PathInt) Group

Rats performed the task in darkness except during the first forced trial each day when a visual cue (white triangle on black background) was briefly presented on the UCM-side video monitor to facilitate orienting to the spatial reference frame at the start of the experiment. After the first trial, there were no visual or tactile sensory cues available to help rats distinguish between the choice points. The first (cued) trial was not included in behavioral scoring. Four unique floor textures (ceramic tile samples, HomeDepot) were placed in the reward zones to distinguish the four corners.

#### Texture Group

Identical to PathInt except that additional, distinct tactile floor textures were placed at the north and south choice points to provide local discriminative cues. Since the room is dark, animals can only access the floor cues when they are physically located at the choice intersections, but not from a distance.

#### Beacon Group

No floor textures were used at choice points nor reward corners. Instead, white LED beacon lights were positioned 5 cm from each UCM reward well and remained illuminated throughout the session. The monitor cue was omitted on the first trial, since the LED beacon provided orienting information throughout the session.

### Phase 2: Revaluation Manipulation

Upon reaching the overtraining criteria, rats underwent 24-hour water deprivation. To enhance thirst during deprivation, 2 g of 8% salted chow was provided during the 25-minute pre-session holding period on the following day. Different cohorts of rats then received one of the following two revaluation manipulations:

#### Incentive Learning (IL)

Rats were confined within the east (room reference frame) segment of the central plus maze by closing adjacent doors. During this session, animals received a two-bottle consumption test in which bottles containing milk and water were suspended from opposite sides of the confinement area, allowing free access to both fluids. No choice points or navigational contingencies were available within this isolated compartment. Thus, rats experienced the altered value of water under deprivation within the maze environment, but without re-experiencing the action–outcome contingencies learned during acquisition. The shift in reward preference was quantified as the proportion of total fluid consumption attributable to milk.

#### Contingency Updating (CU)

Rats performed a full 30-minute acquisition session under water deprivation. Immediately afterward, they received a two-bottle test to confirm a shift of preference from milk to water.

The two procedures differed only in whether rats re-experienced the learned maze action–outcome contingencies after outcome revaluation. This comparison allowed us to determine whether changes in outcome value influenced subsequent navigational choices under the two revaluation procedures. After completing the manipulation session, rats were returned to ad lib water access and then received a standard reminder task session on the following day without deprivation.

### Phase 3: Strategy Probing

Strategy probe involves two sessions: the hydrated (Hyd) session and the Thirsty (Thr) session. The two sessions were counterbalanced in order. Procedures for each group matched their acquisition sessions, except that no rewards were delivered during both probe sessions. Prior to the Thirsty (but not the Hydrated) test session, rats underwent a 24-hour water deprivation. The Thirsty session assessed whether the change in outcome value induced by water deprivation influenced subsequent navigational choice behavior, while the Hydrated session serves as a control. Immediately after each probe session, rats received a two-bottle consumption test to verify reward preference.

### Phase 4: HPC Inactivation (Intracranial Infusion)

Each animal received two infusion sessions: VEH (aCSF, 0.5 µL per hemisphere) and Muscimol (1 µg/µL; 0.25 µL per hemisphere). The order of aCSF and 0.25 µL muscimol sessions was counterbalanced across animals. Cannula placement was verified histologically following completion of behavioral testing (see Histology section).

Bilateral intracranial infusions were administered using 28-gauge internal cannulas extending 1 mm beyond the guide cannula tips (Protech International Inc.). Internal cannulas were connected via PE tubing (BD Intramedic™) to 5 µL Hamilton syringes (Model 75N), which were mounted on a microinfusion pump (Harvard Apparatus). To prevent mixing of drug and oil within the tubing, syringes were backfilled with peanut oil and separated from the drug solution by a small air bubble. The presence and movement of this bubble during infusion allowed visual confirmation of successful drug delivery. Muscimol (1 µg/µL; MilliporeSigma) or artificial cerebrospinal fluid (aCSF; Tocris Bioscience) was loaded into the infusion line immediately prior to use. Drug-containing tubing was kept chilled on ice and shielded from light until infusion.

For infusion, rats were gently restrained and dummy cannulas were removed and replaced with internal cannulas. Infusions were delivered at a rate of 0.25 µL/min. Total infusion volumes were 0.25 µL or 0.5 µL per hemisphere, depending on session. After infusion completion, injectors remained in place for 2 minutes to allow diffusion before being slowly withdrawn and replaced with sterile dummy cannulas. Following infusion, animals were placed in a holding cage under a dark cloth for 45 minutes prior to behavioral testing to allow drug diffusion. Animals were monitored continuously during this period. All infusion equipment was cleaned with 200-proof ethanol after each session.

### Statistical analysis

All analyses of variance (ANOVA) were performed on JASP (JASP Team, 2026). When session type was a within-subject independent variable, the two target sessions of each animal were truncated to the same trial number. Repeated-measure (RM) ANOVAs are followed by planned linear contrasts. Contrast statistics (t value, effect size *d*_z_, the 95% confidence interval of the effect size) are computed using pooled error term, aligning with the RM-ANOVA structure.

### Histology

Rats were anesthetized (isoflurane, VetOne), euthanized (i.p. injection of 1 mL pentobarbital, FATAL-PLUS), and transcardially perfused with 200 mL PBS (0.01 M, Fisher BioReagents™) and 200 mL 4% paraformaldehyde (PFA, Thermo Scientific Chemicals). Brains were extracted and soaked in 30% sucrose PFA for at least seven days before they were sectioned at 40 μm on a cryostat (Leica CM1950). Brain slices were mounted on slides (Fisherbrand^TM^ Superfrost^TM^ Plus) and imaged on a microscope (Keyence BZ-X). Infusion sites extracted from histology were shown on Figure 6C.

## Results

Male (M, n = 20) and female (F, n = 22) Long-Evans rats were trained to perform a navigational decision-making task in which identical turning actions (left or right) led to opposite rewards (milk or water) depending on the spatial location at which the turning decision was made (Figure 1, Movie 1). Prior to training, all but 5 rats (all in the Beacon-CU group) underwent survival surgery to bilaterally implant infusion cannulae in the dorsal hippocampus (see Methods). Four male rats assigned to the most difficult task condition failed to reach criterion after 40 days of training and were excluded from subsequent analyses (see Methods).

### Task Design

Rats were trained on a plus maze located at the center of a 2 × 2 m square track (Figure 1A). The maze was centered in a square experiment room with uniformly black walls, floor, and ceiling with no intramaze or extramaze spatial landmark cues. Automatic doors controlled access to maze arms. Before the first training session, rats were confined within an isolated maze segment where they received a two-bottle preference test (Figure 1B, Movie 2) which confirmed a strong baseline preference for unsweetened condensed milk over water (see below).

During training trials (Figure 1C), two corners (northwest and southwest) were baited with milk (100 μL) and the other two (northeast and southeast) with water (300 μL). To begin each trial (Figure 1C, top), rats approached the maze center from either the west or east arm (randomized by trial). They were then forced to the north or south choice point (independently randomized by trial) where they freely decided to turn left (L) or right (R) (Figure 1C, middle). The turning response for reaching the preferred milk well was reversed between choice locations (north choice point: L→milk, R→water; south choice point: L→water, R→milk). After reaching the chosen well, rats were guided back to the next start arm along one of several forced return paths (Figure 1C, bottom). The return path was randomly selected on each trial.

To prevent use of odor cues, inaccessible decoy milk wells were placed beneath all four corners. Throughout each session, the maze room was flooded with white noise (60 dB at maze center) through a central overhead speaker. Prior to each session, the task reference frame was rotated with respect to the experiment room to minimize uncontrolled extramaze cues, and all track segments were cleaned, removed, and replaced in scrambled order to minimize uncontrolled intramaze cues.

Rats underwent a 2-phase regimen of training on the task, followed by probe tests to test whether their navigational choices were sensitive or insensitive to outcome revaluation, and by HPC inactivation to verify if the HPC is necessary for the navigation task (Figure 2A).

### Phase 1: Acquisition and Overtraining

Rats were assigned to one of three training conditions (Figure 2C): 1) a Beacon Group (6M, 10F) trained with LED lights adjacent to the milk wells, allowing them to approach reward by following a visual beacon without need of distinguishing between North vs. South choice points; 2) a Texture Group (6M, 6F) trained in darkness with distinct floor textures at the two intersections, enabling local tactile, but not remote visual, perceptual discrimination between the North vs. South choice points, 3) a Path Integration (PathInt) Group (4M, 6F) trained in darkness without sensory cues at choice points, requiring rats to rely on memory of their own self-motion (path integration) for discrimination between the North vs. South choice points.

All groups were trained daily for 30 min until they met the following overtraining criteria. On two consecutive days, rats were required to: 1) choose milk on ≥8 of the first 10 trials, 2) choose milk on ≥ 90% of all visits to each choice point during the session, and 3) run ≥ 30 trials per session. Figure 2B shows the number of sessions required to reach this overtraining criterion for rats in each group. A one-way ANOVA revealed a significant effect of training condition (F(2,35) = 33.87, p < .001, η_p_ ^2^ = 0.66). PathInt rats required the most sessions to reach criterion (24.60 ± 2.37), followed by Texture (12.92 ± 1.57) and Beacon (8.13 ± 0.43) rats. Tukey post hoc tests confirmed slower acquisition in PathInt rats than Texture (t(35) = 5.46, p < .001, *d*_z_ = 2.34, 95% CI [1.05, 3.63]) and Beacon (t(35) = 8.18, p < .001, *d*_z_ = 3.30, 95% CI [1.88, 4.72]) rats. Beacon rats learned fastest, significantly faster than Texture rats (t(35) = 2.51, p = 0.04, *d*_z_ = 0.96, 95% CI [-0.04, 1.96]).

#### Sex effect in the PathInt group

Ten rats successfully acquired the PathInt version of the task (4M, 6F), but an additional four male rats failed to reach criterion after 40 days of training and were excluded from the study. Inspection of acquisition rates suggested a marked sex difference in the PathInt condition, with females reaching criterion substantially more rapidly than males (Figure 2B).

A two-way ANOVA with training group and sex as factors confirmed both a main effect of training group (F(2,32) = 54.07, p < .001, ηp² = 0.77) and a main effect of sex (F(1,32) = 12.18, p = 0.001, ηp² = 0.28), qualified by a significant sex × group interaction (F(2,32) = 5.32, p = 0.01, ηp² = 0.25). Uncorrected post hoc comparisons revealed no significant sex difference in the Texture (t(32) = 0.77, p = 0.45, *d*_z_ = 0.45, 95% CI [-0.74, 1.63]) or Beacon (t(32) = 0.53, p = 0.60, *d*_z_ = 0.28, 95% CI [-0.78, 1.33]) groups. In contrast, females acquired the PathInt task substantially faster than males (t(32) = 4.33, p < .001, *d*_z_ = 2.80, 95% CI [1.30, 4.29]).

Because sex differences were not a primary focus of the present study, we did not investigate this effect further. If replicable, the pronounced female advantage observed in the PathInt condition suggests that this version of the task may provide a useful assay for future investigations of sex differences in spatial learning and memory.

### Phase 2: Incentive Learning (IL) or Contingency Updating (CU)

After reaching criterion at the end of Phase 1, rats in each training group were split into two cohorts for Phase 2 (Figure 3A): Incentive Learning (IL: 10F, 8M) and Contingency Updating (CU: 12F, 8M) (Table 1). IL and CU cohorts both underwent 24 hours of water restriction at the beginning of Phase 2 (Figure 3B), after which their regimens differed.

**Figure 3.**
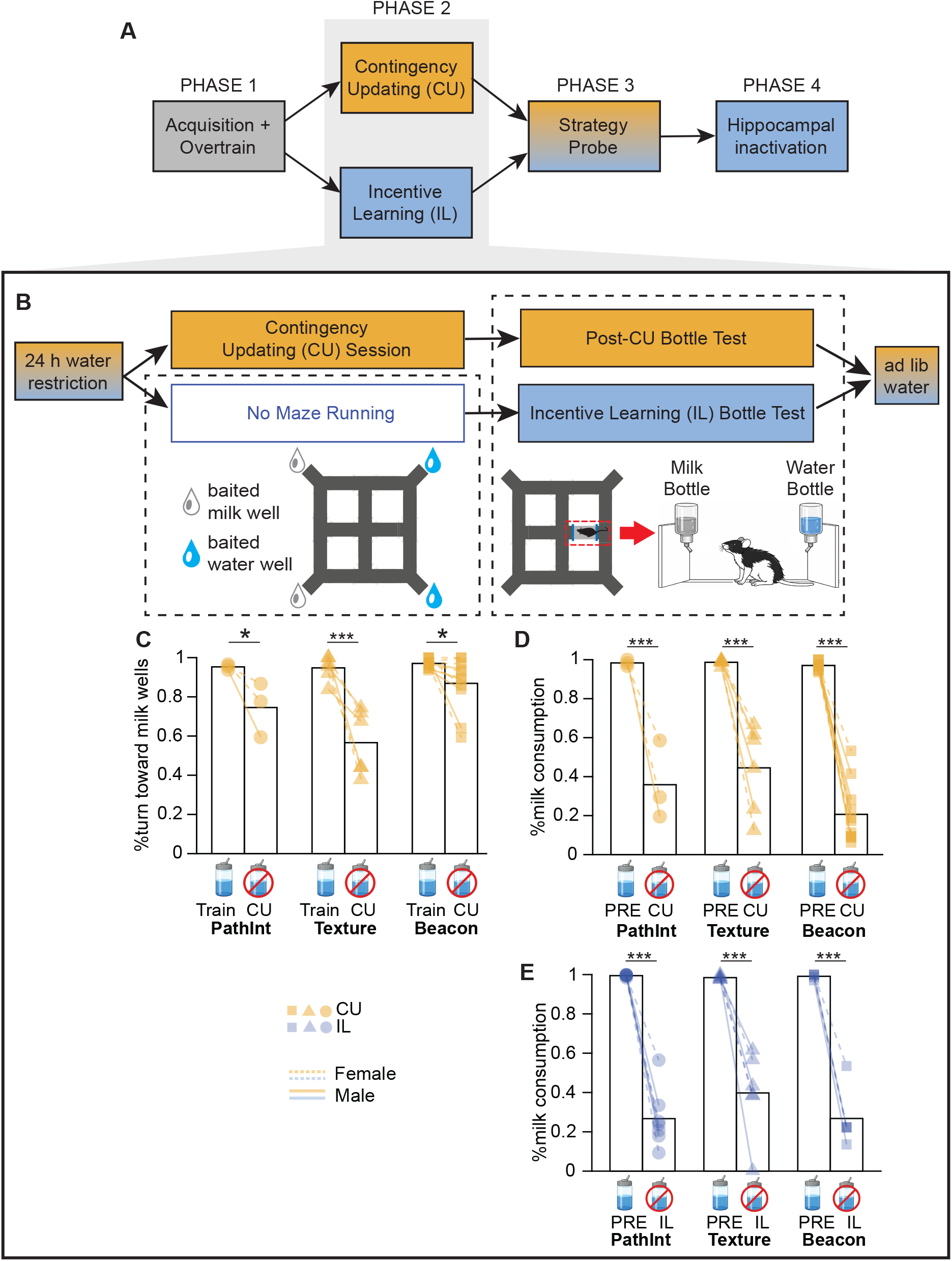
Revaluation Training and Incentive Learning A) This figure shows results from Phase 2 of the experiment, given after the overtraining criteria had been reached in Phase 1. B) Timeline for phase 2: After 24-hr water restriction, animals in each of the three training groups were split into two cohorts, one receiving Revaluation Training (CU; followed immediately by a bottle test) and the other receiving a bottle test only (Incentive Learning, IL) without maze training; all rats were returned to ad lib water after the session. C) CU cohort’s choice behavior on the baited maze (% turning choices toward milk wells, y-axis) is compared during hydrated (final acquisition session, Train) vs. thirsty (CU session) conditions for the PathInt-CU, Texture-CU, and Beacon-CU groups. D) CU cohort’s milk consumption on bottle tests (milk volume as % all fluid consumed, y-axis) is compared for hydrated (pre-training bottle test, PRE) vs. thirsty (CU) conditions in all three CU groups. E) IL cohort’s milk consumption on bottle tests is compared for hydrated (pre-training bottle test, PRE) vs. thirsty (IL) conditions in all three IL groups. Planned comparisons: *p < .05, ***p < .001.

#### CU cohort

The CU cohort included 20 rats: 3 PathInt rats (2F, 1M), 6 Texture rats (3F, 3M), and 11 Beacon rats (7F, 4M). Water-restricted rats in the CU cohort were given a standard training session on the maze while thirsty (Figure 3B). This regimen allowed CU rats to experience the altered relative values of milk and water under thirst while also re-experiencing the learned maze action–outcome contingencies in this altered motivational state, thereby providing an opportunity for those contingencies to be updated.

Figure 3C shows that CU rats reduced milk-associated responding during the CU session relative to the final training session. A 2 × 3 mixed ANOVA with session (Train vs. CU) as a repeated factor and training group (PathInt, Texture, Beacon) as a between-subjects factor revealed a significant main effect of session, indicating a shift away from milk-associated responses under thirst (F(1,17) = 34.57, p < .001, η_p_ ^2^ = 0.67). The session × group interaction was also significant (F(2,17) = 6.52, p = 0.008, η_p_ ^2^ = 0.43), indicating that the magnitude of this shift differed among groups. These large omnibus effects indicate that the CU manipulation produced a robust overall shift in responding from milk-to water-associated responses, and that this shift varied substantially across training groups.

Follow-up planned contrasts for simple effects showed reductions in milk-associated responding in all three groups in the predicted direction (PathInt-CU: t(17) = 2.35, p = 0.03, *d*_z_ = 1.96, 95% CI [0.13, 3.79]; Texture-CU: t(17) = 6.12, p < .001, *d*_z_ = 3.61, 95% CI [1.87, 5.34]; Beacon-CU: t(17) = 2.21, p = 0.04, *d*_z_ = 0.96, 95% CI [0.01, 1.91]; two-tailed). The PathInt-CU effect should be interpreted cautiously because the subgroup sample size was small (n=3) and the wide *d*_z_ confidence interval indicates low precision in the estimated effect size. The *d*_z_ confidence interval was also relatively wide for the Beacon-CU effect despite this being the largest subgroup (n = 11), suggesting substantial variability in how Beacon-CU rats adjusted their responding under deprivation. Notably, Beacon-CU showed the smallest mean reduction in milk-associated responding, suggesting that Beacon-CU rats may have been less influenced by the revaluation manipulation than rats in the other training conditions.

Immediately after the CU session, rats were given a two-bottle test in a confined segment of the maze. Figure 3D shows that all three training groups consumed less milk (measured as % of total fluid consumed) during the Post-CU bottle test (given under water restriction) than during their original preference test given prior to acquisition training. A mixed 2×3 ANOVA found a significant main effect of session (PRE vs. CU: F(1,17) = 199.83, p <. 001, η_p_ ^2^ = 0.92) indicating a robust shift in preference from milk to water under thirst. There was also a significant session × group interaction effect (F(2,17) = 3.95, p = 0.04, η_p_ ^2^ = 0.32), possibly because Beacon-CU rats were thirstier than other groups during the two-bottle test after consuming less water during the prior maze session (because of their greater perseverance on milk-associated responses).

#### IL cohort

There were n=18 rats in the IL cohort: 7 (4F, 3M) in the PathInt group, 6 (3F, 3M) in the Texture group, and 5 (3F, 2M) in the Beacon group. Water-restricted rats in the IL cohort were confined within an isolated maze segment where they received a two-bottle preference test (Figure 3B), allowing them to experience the altered relative values of milk and water under thirst (incentive learning). However, IL rats were not challenged to make decisions at choice points on the maze in their thirsty state, and therefore did not re-experience the learned action– outcome contingencies under thirst as the CU cohort did.

Figure 3E shows that milk intake during the two-bottle test dropped sharply after water restriction (IL session) compared with the two-bottle test administered prior to acquisition training when rats had free access to water in the homecage (PRE session; see Figure 1B). A 2 × 3 mixed ANOVA with session (PRE vs. IL) as a repeated factor and training group (PathInt vs. Texture vs. Beacon) as a between-subjects factor revealed a significant main effect of session (F(1,15) = 267.23, p < .001, η_p_ ^2^ = 0.95), indicating a robust shift in preference from milk to water under thirst. Follow-up within-group planned contrasts confirmed large reductions in milk preference in all three subgroups (PathInt-IL: t(15) = 11.02, p < .001, *d*_z_ = 5.83, 95% CI [3.40, 8.26]; Texture-IL: t(15) = 8.23, p <. 001, *d*_z_ = 4.71, 95% CI [2.60, 6.81]; Beacon-IL: t(15) = 9.26, p <. 001, *d*_z_ = 5.80, 95% CI [3.29, 8.31]).

In summary, Phase 2 results showed that both CU and IL cohorts experienced robust shifts in outcome preference following water deprivation. The critical difference between cohorts was that CU animals re-experienced the learned maze contingencies after this change in outcome value, whereas IL animals did not.

### Phase 3: Strategy Probe

At the conclusion of Phase 2, rats were returned to ad lib water overnight. On the following day, they received a reminder training session, followed on subsequent days by two non-rewarded probe sessions: Hydrated (Hyd) and Thirsty (Thr), each immediately followed by a two-bottle preference test. The Hyd and Thr probe sessions were administered two days apart in counterbalanced order, with a standard reminder session on the intervening day (Figure 4A,B).

**Figure 4.**
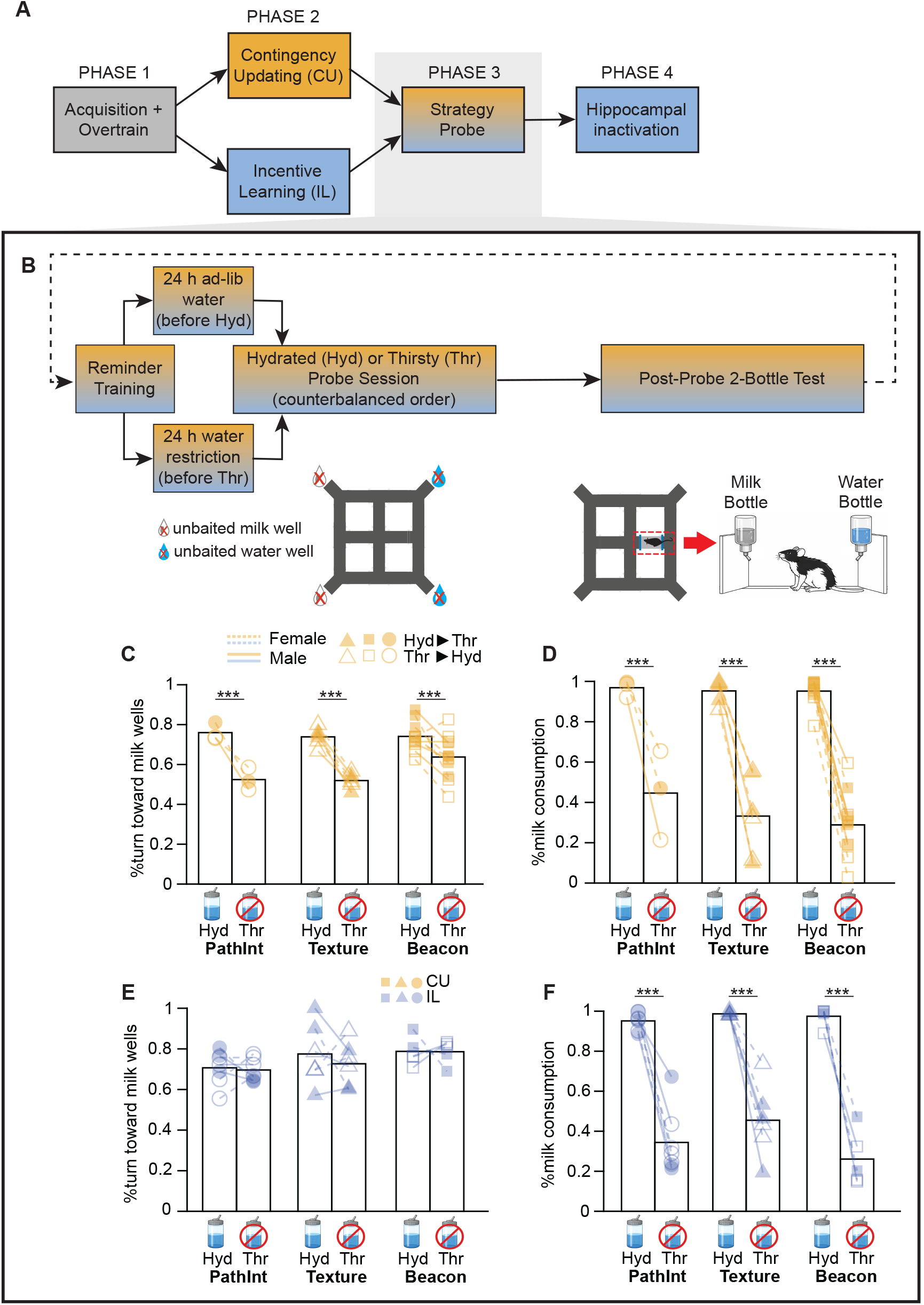
Strategy probe A) This figure shows results from Phase 3 of the experiment. B) Rats received two probe sessions in extinction (that is, on an unbaited maze) in counterbalanced order: Thirsty (Thr, preceded by a reminder session then 24 h water restriction) and Hydrated (Hyd, preceded by a reminder session then ad lib water); immediately after each probe session the rat received a bottle test. C) CU cohort’s choice behavior (% turning choices toward unbaited milk wells, y-axis) during unbaited Thr and Hyd probe sessions. D) CU cohort’s milk consumption on bottle tests (milk volume as % all fluid consumed, y-axis) during Thr vs. Hyd conditions. E) IL cohort’s choice behavior during unbaited Thr and Hyd probe sessions. F) IL cohort’s milk consumption on bottle tests during Thr and Hyd probe sessions. Planned comparisons: ***p < .001.

#### CU cohort

Figure 4C shows that thirsty CU rats shifted their navigational choices away from milk-associated responses during non-rewarded probe sessions in all three CU subgroups (PathInt-CU, Texture-CU, and Beacon-CU). A 2 × 3 mixed ANOVA with session (Hyd vs. Thr) as a repeated factor and training group (PathInt, Texture, Beacon) as a between-subjects factor revealed a main effect of session (F(1,17) = 65.26, p <. 001, η_p_ ^2^ = 0.79) with no main effect of group (F(2,17) = 2.03, p = 0.16, η_p_ ^2^ = 0.19). Although there was a significant session × group interaction (F(2,17) = 4.55, p = 0.03, η_p_ ^2^ = 0.35), follow-up planned contrasts showed robust reductions in milk-associated responding in all three groups in the predicted direction (PathInt: t(17) = 4.54, p < .001, *d*_z_ = 3.06, 95% CI [1.32, 4.80]; Texture: t(17) = 5.98, p < .001, *d*_z_ = 2.85, 95% CI [1.46, 4.25]; Beacon: t(17) = 3.81, p = 0.001, *d*_z_ = 1.34, 95% CI [0.48, 2.21]; two-tailed).

Two-bottle preference tests conducted immediately after the probe sessions confirmed a robust revaluation effect, demonstrating a preference shift toward water in thirsty rats (Figure 4D). Water deprivation significantly reduced milk preference (main effect of session: F(1,17) = 278.87, p < .001, η_p_ ^2^ = 0.94) with no main effect of group (F(2,17) = 0.71, p = 0.51, η_p_ ^2^ = 0.08) and no session × group interaction (F(2,17) = 1.20, p = 0.33, η_p_ ^2^ = 0.12). Thus, CU rats showed clear outcome revaluation during bottle testing, and their navigational choices also shifted following revaluation. The present experiments do not distinguish whether this shift reflected a transition to goal-directed control based on anticipation of future outcomes, or instead reflected another mechanism, such as acquisition of state-dependent stimulus-response associations during contingency updating (see Discussion). In either case, the observed response shift in the CU cohort provides a positive control demonstrating that the task design was capable of detecting motivationally-induced changes in navigational choice behavior.

#### IL cohort

Figure 4E shows that choice behavior during non-rewarded probe sessions showed little evidence of revaluation sensitivity in the IL subgroups (PathInt-IL, Texture-IL, and Beacon-IL) . A 2 × 3 mixed ANOVA with session (Hyd vs. Thr) as a repeated factor and training group (PathInt, Texture, Beacon) as a between-subjects factor revealed no main effect of session (F(1,15) = 0.36, p = 0.56, η_p_ ^2^ = 0.023), no main effect of group (F(2,15) = 2.61, p = 0.11, η_p_ ^2^ = 0.26), and no session × group interaction (F(2,15) = 0.18, p = 0.84, η_p_ ^2^ = 0.02). The absence of both a session effect and a session × group interaction indicates that water deprivation did not reliably shift rats from milk-associated toward water-associated responses in any IL training condition. Follow-up planned contrasts likewise showed only small and nonsignificant changes in milk-associated turning responses (PathInt-IL: t(15) = 0.20, p = 0.84, *d*_z_ = 0.11, 95% CI [-1.03, 1.25]; Texture-IL: t(15) = 0.84, p = 0.42, *d*_z_ = 0.50, 95% CI [-0.74, 1.74]; Beacon-IL: t(15) = 0.03, p = 0.98, *d*_z_ = 0.02, 95% CI [-1.33, 1.36]; two-tailed). Direct comparison of CU and IL cohorts (reported below) confirmed that revaluation effects were significantly larger following CU than IL training.

In contrast with this lack of a revaluation effect on choice behavior, two-bottle preference tests conducted immediately after the probe sessions confirmed a robust revaluation effect upon consumption (Figure 4F). Water deprivation significantly increased water consumption relative to milk consumption (main effect of session: F(1,15) = 254.65, p < .001, η_p_ ^2^ = 0.94), with no main effect of group (F(2,15) = 2.18, p = 0.15, η_p_ ^2^ = 0.23) and no session × group interaction (F(2,15) = 1.71, p = 0.21, η_p_ ^2^ = 0.19). Hence, although IL rats showed clear reward revaluation during bottle testing, they showed little evidence of incorporating this updated outcome value into navigational choices on the maze, consistent with habitual or outcome-insensitive behavioral control.

#### Comparing revaluation in IL vs. CU cohorts

To directly compare the effects of contingency updating on subsequent choice behavior, we computed a Revaluation Index (RI) for each rat: RI = %M_Hyd_ - %M_Thr_, where %M_Hyd_ and %M_Thr_ represent the percentage of turning responses directed toward unbaited milk wells during hydrated and thirsty probe sessions, respectively (Figure 5). Positive RI values indicate that thirst shifted responding away from milk-associated choices and toward water-associated choices, whereas RI values near zero indicate little or no effect of thirst on choice behavior.

**Figure 5.**
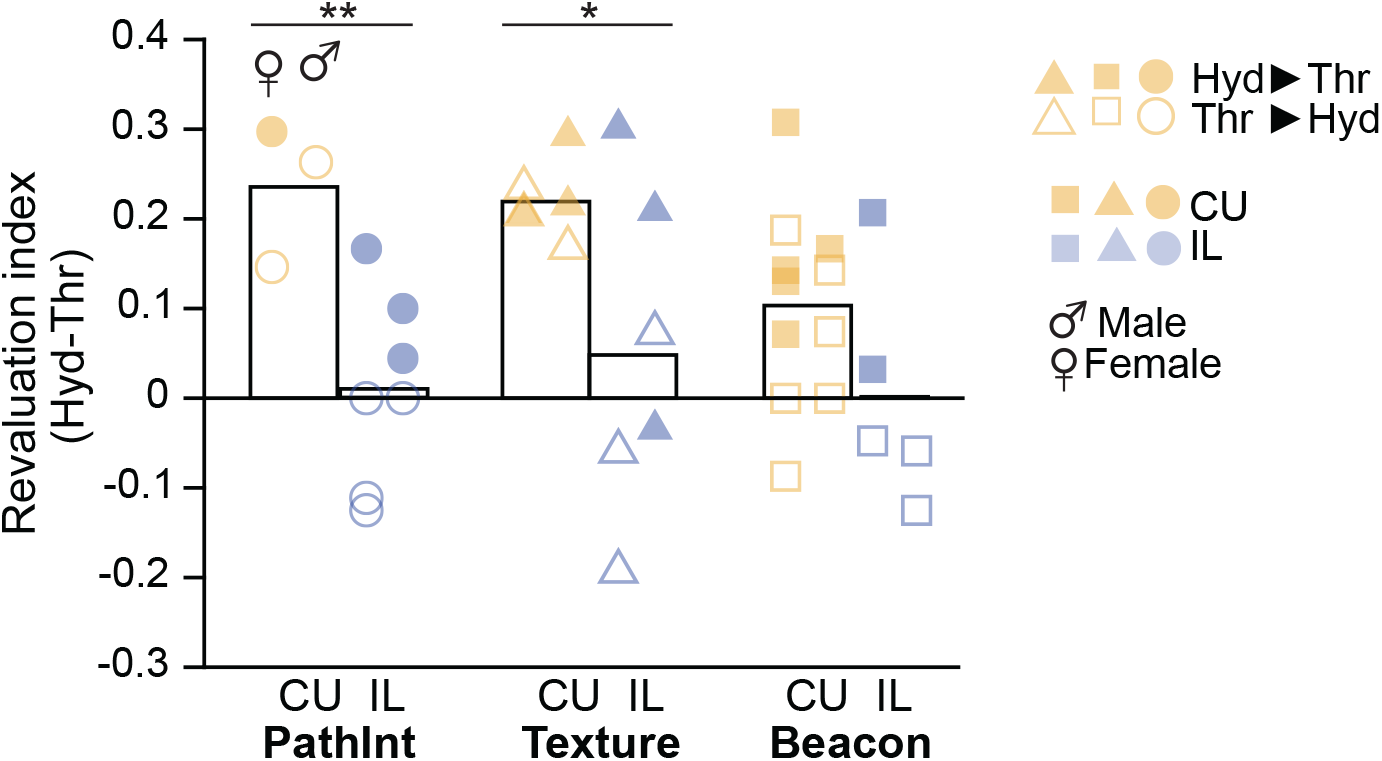
Revaluation effect in CU versus IL cohort Bars plot the mean revaluation index (y-axis; % turns toward milk when hydrated minus when thirsty) from strategy probe sessions for rats in CU versus IL cohorts within each training group (in scatter plots, points for male and female rats are plotted to the left and right side, respectively, of each bar). Asterisks show results from planned CU vs IL contrasts; *p < .05, **p < .01.

A 2 × 3 between-subjects ANOVA on RI values with cohort (CU vs. IL) and training group (PathInt, Texture, Beacon) as factors revealed a significant main effect of cohort (CU vs IL: F(1,32) = 16.49, p < .001, η_p_ ^2^ = 0.34), but neither a main effect of training group (F(2,32) = 1.80, p = 0.18, η_p_ ^2^ = 0.10) nor a cohort × group interaction (F(2,32) = 0.76, p = 0.48, η_p_ ^2^ = 0.05). Thus, CU rats that re-experienced the maze contingencies under thirst exhibited significantly larger shifts in navigational choice behavior than IL rats that experienced incentive learning alone, and this effect was broadly similar across training conditions. Planned comparisons between CU and IL cohorts further supported this conclusion. Revaluation indices were significantly larger in CU than IL rats in both the PathInt (t(32) = 2.79, p = 0.01, *d*_z_ = 1.92, 95% CI [0.44, 3.41]) and Texture (t(32) = 2.54, p = 0.02, *d*_z_ = 1.47, 95% CI [0.23, 2.70]) groups, indicating that thirst shifted navigational choices toward water-associated responses after contingency updating but not after incentive learning alone. The corresponding CU vs IL comparison was not significant in the Beacon group (t(32) = 1.62, p = 0.12, *d*_z_ = 0.87, 95% CI [-0.25, 1.99]), and this was accounted for by an absence of response shifting in Beacon-CU rats rather than a presence of response shifting in Beacon-IL rats, reflecting the comparatively smaller revaluation effect observed in Beacon-CU rats.

Together, these analyses demonstrate that explicit contingency updating produced substantially larger changes in navigational choice than incentive learning alone. Because both cohorts exhibited comparably robust shifts in outcome preference during bottle testing, this difference cannot readily be attributed to differences in the effectiveness of the revaluation manipulation itself. Rather, the IL cohort exhibited outcome-insensitive navigational choices despite clear evidence that outcome values had changed, consistent with the widely used operational criterion for distinguishing habitual from goal-directed control based on sensitivity to outcome revaluation.

### Phase 4: Hippocampal Inactivation

To assess hippocampal contributions to choice behavior, rats from the IL cohort received intracranial infusions of vehicle (VEH, 0.5 µL aCSF) and muscimol (MUS, 0.25 µL per hemisphere at 1 µg/µL) in counterbalanced order during standard maze sessions (Figure 6A,B). Each infusion session (VEH or MUS) was preceded by a drug-free reminder session. Figure 6C shows injection sites superimposed on an atlas plate from Swanson (2018). Three rats from the PathInt-IL group were excluded after postmortem determination that guide cannulae of incorrect length had been implanted, reducing the PathInt-IL sample size to (n=4) for drug infusion analyses (Table 2). Similar to the full PathInt-IL group, the infused PathInt rats were insensitive to outcome revaluation on the maze (Figure 6E; t(3) = 0.10, p = 0.93, *d*_z_ = 0.05, 95% CI for *d*_z_ [-0.93, 1.03], paired t-test), but not in the bottle test (Figure 6F; t(3) = 5.85, p = 0.01, *d*_z_ = 2.92, 95% CI for *d*_z_ [0.51, 5.34], paired t-test).

**Figure 6.**
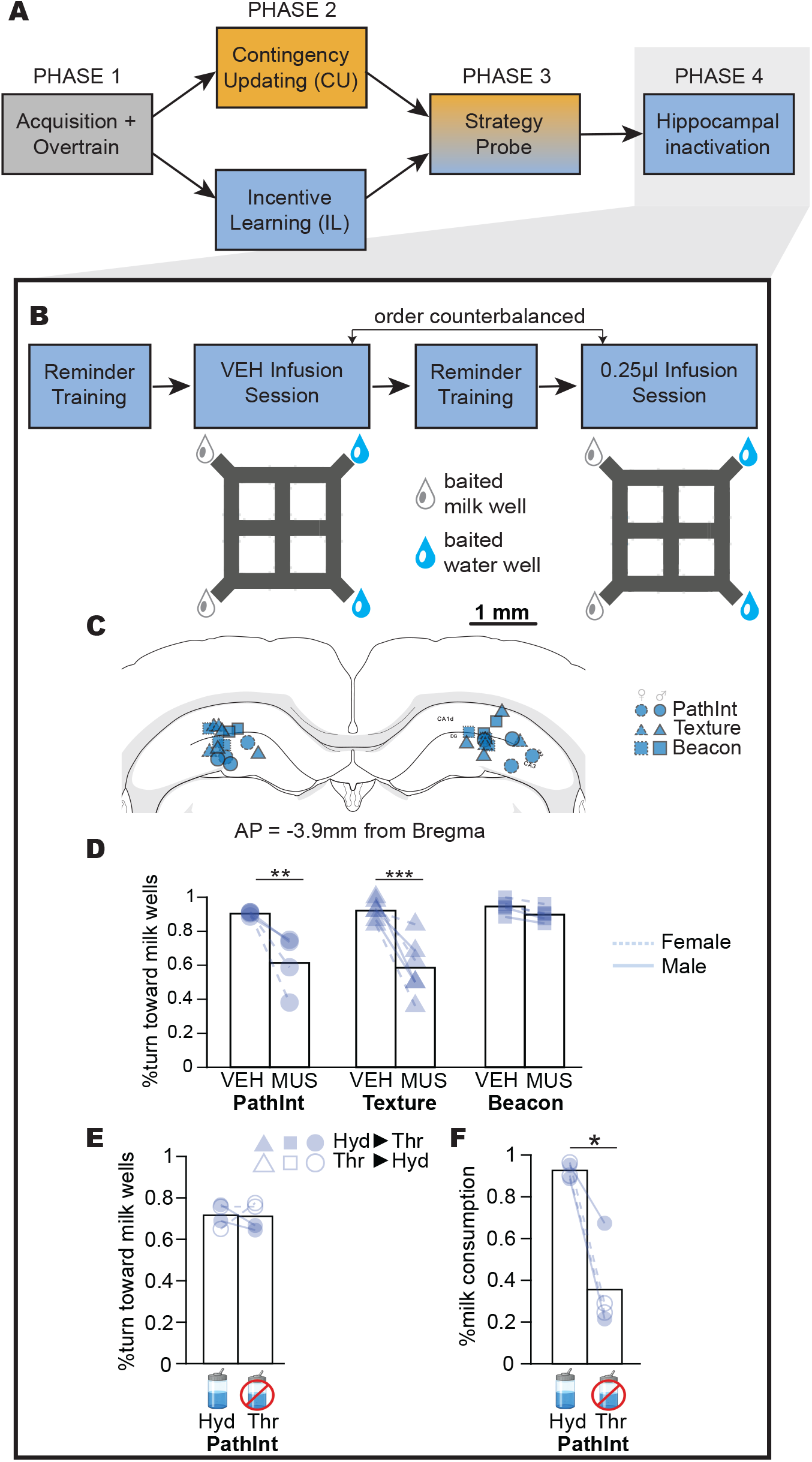
HPC inactivation A) This figure shows results from Phase 4 of the experiment. B) Timeline of phase 4: IL animals received separate infusions of vehicle (VEH) and muscimol (MUS) in counterbalanced order, interleaved with regular reminder training sessions. C) Histological reconstruction of Infusion sites in dorsal hippocampus. D) IL cohort’s choice behavior (% turning choices toward milk wells, y-axis) during standard sessions after infusions of VEH or MUS. E,F) Strategy probe (panel E) and bottle test (panel F) results are plotted separately for the subset of four PathInt-IL animals that were included in the infusion analysis. Planned comparisons: *p < .05, **p < .01, ***p < .001.

Choice behavior after infusion is shown in Figure 6D. A 2 × 3 mixed ANOVA with training group (PathInt-IL, Texture-IL, Beacon-IL) as a between-subjects factor and drug condition (VEH vs. MUS) as a repeated factor revealed a significant group × drug interaction (F(2,12) = 6.85, p = 0.01, η_p_ ^2^ = 0.53). Follow-up planned contrasts revealed significant reductions in choosing the milk-associated response following muscimol infusion in the PathInt-IL (t(12) = 4.31, p = 0.001, *d*_z_ = 2.76, 95% CI for *d*_z_ [1.00, 4.53]) and Texture-IL (t(12) = 6.12, p < .001, *d*_z_ = 3.20, 95% CI for *d*_z_ [1.47, 4.94]) groups, but not in the Beacon-IL group (t(12) = 0.79, p = 0.45, *d*_z_ = 0.45, 95% CI for *d*_z_ [-0.75, 1.66]). Although the Beacon-IL confidence interval was relatively wide, the observed effect size was substantially smaller than those observed in the PathInt-IL and Texture-IL groups. Thus, hippocampal inactivation impaired successful navigation toward the milk-associated well when choosing the milk-associated response required discrimination between spatially defined choice locations, but such impairment was not seen when navigation could be guided by visual beacons.

Planned interaction contrasts further confirmed that muscimol impairment was significantly greater in the hippocampal-dependent conditions than in the beacon-guided condition (PathInt-IL vs. Beacon-IL: t(12) = 2.68, p = 0.02, *d*_z_ = 2.31, 95% CI [0.28, 4.34]; Texture-IL vs. Beacon-IL: t(12) = 3.54, p = 0.004, *d*_z_ = 2.75, 95% CI for *d*_z_ [0.77, 4.74]). These contrasts indicate that the absence of a reliable muscimol effect in the Beacon-IL group was not merely attributable to reduced statistical power, but instead reflected a substantially smaller impairment than that observed in task variants requiring hippocampal-dependent discrimination between spatially defined choice locations.

Under vehicle infusion, all three groups selected the milk-associated response significantly more often than expected by chance (PathInt: t(3) = 66.69, p < .001, *d*_z_ = 33.34, 95% CI [11.40, ∞]; Texture: t(5) = 16.62, p < .001, *d*_z_ = 6.79, 95% CI [3.18, ∞]; Beacon: t(4) = 23.88, p < .001, *d*_z_ = 10.68, 95% CI [4.45, ∞]; Student’s t test against 0.5). Following muscimol infusion, however, performance in the PathInt-IL and Texture-IL groups no longer significantly exceeded chance (PathInt: t(3) = 1.33, p = 0.14, *d*_z_ = 0.66, 95% CI [-0.31, ∞]; Texture: t(5) = 1.24, p = 0.14, *d*_z_ = 0.51, 95% CI [-0.24, ∞]), whereas Beacon-IL performance remained significantly above chance (t(4) = 21.04, p < .001, *d*_z_ = 9.41, 95% CI [3.91, ∞]). Thus, above-chance selection of milk associated responses was no longer detectable following hippocampal inactivation.

Taken together, these findings demonstrate that accurate performance in the PathInt-IL and Texture-IL task variants depended upon intact dorsal hippocampal function. Temporary hippocampal inactivation selectively disrupted navigational performance in these groups, eliminating reliable above-chance selection of the preferred milk-associated response, whereas performance in the Beacon-IL group remained largely unaffected. Importantly, the same PathInt-IL and Texture-IL groups had previously exhibited outcome-insensitive navigational choices during the strategy probe (Figures 4E and 5E). Thus, successful navigation in these task variants required hippocampal processing even though navigational choices remained insensitive to outcome revaluation. To our knowledge, these findings provide the first direct evidence that hippocampal-dependent navigation can coexist with outcome-insensitive choice behavior.

## Discussion

The hippocampus is widely implicated in spatial navigation and model-based behavior supported by cognitive maps (O’Keefe and Nadel, 1978; Olton and Samuelson, 1976; Olton et al., 1982; Morris et al., 1982; Packard and McGaugh, 1992, 1996). Prior studies have implicated the hippocampus in goal-directed, outcome-sensitive decision making (Goodman et al., 2016; Miller et al., 2017; Kosaki et al., 2018; Bradfield et al., 2020). However, few studies have examined whether it can also support outcome-insensitive action selection, widely regarded as a behavioral criterion for habitual responding.

Here we present novel evidence that outcome-insensitive (habitual) action selection can occur during a navigation task that depends on the dorsal hippocampus. Hippocampal inactivation impaired navigation in the PathInt-IL and Texture-IL groups, but not in the Beacon-IL group. Outcome-insensitive choice behavior nevertheless occurred in all groups, including the hippocampal-dependent groups. Thus, by demonstrating habitual responding under conditions in which accurate navigation depended upon the dorsal hippocampus, our findings provide evidence that hippocampal representations may support habitual action selection, but alternative interpretations are also possible.

### Mnemonic, predictive, and evaluative roles of the cognitive map

The present findings indicate that hippocampal-dependent navigation can occur in the absence of outcome sensitivity. PathInt-IL and Texture-IL rats depended upon the dorsal hippocampus for accurate navigation to the milk well when milk was the preferred reward, yet they continued to select milk-associated responses after their preference shifted from milk to water, consistent with habitual action selection. Thus, hippocampal dependence on navigation did not imply sensitivity to current outcome value.

#### Memory without prediction

The ability to predict future outcomes implies the existence of an internal model that generates such predictions. Therefore, decisions that depend upon predictions of future outcomes are generally regarded as model-based decisions. However, internal models may serve functions other than prospective prediction. Reinforcement learning theory holds that model-free decisions are guided by cached action values associated with representations of the current environmental state rather than anticipated future states (Sutton and Barto, 2018). To the extent that a cognitive map is required to infer the current state of the environment (for example, when a rat infers its present spatial location via path integration), and to the extent that cached action values depend upon that state representation, retrospective internal models may contribute to habitual action selection without being used to prospectively predict future outcomes. Under this view, an internal model (cognitive map) can support habitual behavior by enabling accurate retrospective inference of the animal’s current state, rather than by enabling prospective predictions of future states and outcomes as during goal-directed decisions.

This memory-without-prediction view of cognitive map function suggests one possible interpretation of our finding that, although PathInt-IL and Texture-IL rats depended upon the dorsal hippocampus for accurate navigation, their choices remained insensitive to outcome revaluation. Under this interpretation, PathInt-IL and Texture-IL rats relied on hippocampal representations only to infer the current spatial state upon which cached action values depended, without using the hippocampus to predict future states or outcomes. Muscimol therefore disrupted navigation toward the preferred milk wells because rats could no longer discriminate between the two choice locations, preventing retrieval of state-dependent action values upon which habitual choices depended.

#### Prediction without value

An alternative explanation of our present findings is derivable from an alternative view of cognitive map function, which may be termed prediction-without-value. Sensitivity to outcome revaluation implies that a decision is not only model-based but more specifically that it is goal-directed, which can be viewed as a narrower category of model-based decision making. A goal-directed decision may be operationally defined as a choice based upon predictions of future outcomes whose current values reflect the organism’s motivational and physiological conditions at the time of choice (Balleine and Dickinson, 1998). Thus, model-based prediction and outcome-sensitive evaluation need not be identical processes. Rats might rely upon hippocampal predictions of future states to guide navigation, yet still fail to exhibit sensitivity to outcome revaluation if those predictions are not coupled to a value system capable of dynamically incorporating changes in motivational state. Such a possibility is consistent with growing evidence from human neuroscience that the hippocampus learns predictive temporal structure and statistical regularities independently of explicit reward or planning (Schapiro et al., 2016; Turk-Browne, 2019).

This prediction-without-value view of cognitive map function suggests an alternative interpretation of our present findings, namely, that PathInt-IL and Texture-IL rats may have used hippocampal representations to predict future states (such as arrival at a previously rewarded location) while nevertheless failing to update the current value of those predicted states following outcome revaluation. Similar to the conditioned stimulus in Pavlovian sign-tracking (Robinson and Flagel, 2009; Morrison et al., 2015), extensive training may cause spatial states to acquire incentive value that becomes partially detached from the reward itself, or alternatively may cause predictions of reward to remain tied to values learned during training rather than values implied by the animal’s current motivational state. In either case, outcome revaluation would not alter the prediction that a learned action leads to a highly valued location or outcome. Therefore, the revaluation would not alter choice behavior, even if the reward from which that value was originally derived is no longer desired.

### Emergence of Outcome Sensitivity After Contingency Updating

CU cohorts that received explicit revaluation training after acquisition exhibited behavioral sensitivity to outcome revaluation during subsequent strategy probes. The underlying mechanisms responsible for this behavioral sensitivity remain open to interpretation. Under the memory-without-prediction and prediction-without-value frameworks, the observed behavioral sensitivity can be accounted for in different ways.

#### Conjunctive state representations

Under the memory-without-prediction interpretation, revaluation training may have altered representations of the current task state rather than promoting prospective evaluation of future outcomes. For example, during revaluation training, rats may have formed conjunctive representations of spatial location and motivational state (e.g., thirsty versus non-thirsty at the north choice point) and then acquired distinct state-action associations for thirsty versus non-thirsty conditions. Decisions that subsequently depended upon these new state representations would have appeared outcome-sensitive according to conventional operational criteria, even though action selection still relied upon retrieval of cached responses rather than prospective evaluation of future outcomes.

#### Novelty-induced strategy shifts

Alternatively, under the prediction-without-value interpretation, revaluation training may have produced outcome sensitivity in at least two ways. One possibility is that revaluation training introduced novelty by exposing overtrained animals to altered reward values. This novelty may have increased attention to the task and thereby enhanced behavioral sensitivity to outcome value during subsequent decisions, producing sensitivity to outcome revaluation during later strategy probes (Handel and Smith, 2023; Linnebank et al., 2018; Thrailkill et al., 2018; Bouton, 2021).

#### Motivation-sensitive value representations

A second possibility is that revaluation training strengthened the linkage between predicted states and representations of the current value of expected outcomes. Whether dorsal hippocampal neurons encode reward-related signals in addition to spatial states remains an open question that continues to be debated in the literature (Burgess and O’Keefe, 1996; Poucet et al., 2004; Lee et al., 2012; van der Meer et al., 2012; Blanquat et al., 2013; Gauthier and Tank, 2018; Duvelle et al., 2019; Hampson and Deadwyler, 2000; Masuda et al., 2020). If reward-related signals do exist in the dorsal hippocampus, they may remain tied to values acquired during training rather than the current value of the outcome following revaluation. Such a dissociation could help explain why the PathInt-IL and Texture-IL groups, despite depending upon the hippocampus for accurate navigation, failed to exhibit outcome sensitivity. Revaluation training may have enabled rats to make decisions based upon the current value (at the time of choice) of expected outcomes, either by allowing hippocampal representations to reflect the current value of expected outcomes following revaluation or by strengthening associations between hippocampal state representations and other brain regions that encode outcome values in a motivation-dependent manner.

One way to test this possibility would be to determine whether hippocampal activity is required during acquisition of outcome-sensitive control rather than during expression of outcome-sensitive behavior. For example, if dorsal hippocampal inactivation during revaluation training, but not during revaluation testing, were to impair subsequent outcome sensitivity in Beacon-CU rats (which can perform the standard navigation task independently of the hippocampus), this would suggest that the hippocampus contributes to acquisition of value-sensitive control independently of its role in spatial navigation.

## Conclusions

In summary, our results provide novel evidence that the hippocampus can support navigational decisions that are insensitive to the current value of rewards, a widely used operational criterion for habitual control. Thus, reliance upon a hippocampal cognitive map does not necessarily imply outcome-sensitive, goal-directed decision making during spatial navigation. Further work is needed to distinguish whether hippocampal representations contributed primarily to state inference, prospective state prediction, or motivation-sensitive evaluation. By allowing hippocampal dependence and outcome sensitivity to be manipulated independently, the present task provides a useful approach for addressing these questions and clarifying how hippocampal representations interact with motivational and valuation systems during transitions between habitual and goal-directed control.

## Movie captions

Movie 1. Trial structure and group-specific variants (overhead view)

Section 1.1) Syringe pumps in the two west and two east corners contain unsweetened condensed milk and drinking water, respectively. The rat starts from the west or the east start arm, and makes turning decisions at the north and south choice points. The room is illuminated with invisible IR light and is flooded with white noise. The PathInt group has four unique floor textures at the reward corners but no floor textures at choice points. Section 1.2) The Texture group has two unique floor textures at the choice points, in addition to the textures at the reward corners. Section 1.3) The Beacon group has a LED light next to each milk reward, and has no floor textures in any part of the maze.

Movie 2. Two-bottle preference test structure (side view)

The rat is confined to the east start arm and the room is illuminated with invisible IR lights and is flooded with white noise. The rat has access to two bottles, containing 75ml of unsweetened condensed milk or drinking water. Start location of the bottles randomizes across sessions. The bottles are swapped every 4min. Weight of the bottles are recorded before and after the preference test.

## Acknowledgements

This work was supported by NIH U01NS128664 and NIH U01NS126050 to H.T.B. We thank Asal Esteghlal, Elena Chen, Cristal Cruz, and Lorelei Mae Cordes for assistance with data collection. We thank Kate Wassum, Alicia Izquierdo, Melissa Sharpe, and Andrew Wikenheiser for helpful comments on the experimental design.

